# A feed-forward loop between SorLA and HER3 determines heregulin response and neratinib resistance

**DOI:** 10.1101/2020.06.10.143735

**Authors:** Hussein Al-Akhrass, James R.W. Conway, Annemarie Svane Aavild Poulsen, Ilkka Paatero, Jasmin Kaivola, Artur Padzik, Olav M. Andersen, Johanna Ivaska

## Abstract

Current evidence indicates that resistance to HER2-targeted therapies is frequently associated with HER3 and active signaling *via* HER2-HER3 dimers, particularly in the context of breast cancer. Thus, understanding the response to HER2-HER3 signaling and the regulation of the dimer *per se* remains essential to decipher therapy relapse mechanisms. Here, we demonstrate that signaling by HER3 growth factor ligands, heregulins, support the transcription of a type-1 transmembrane sorting receptor, sortilin-related receptor (SorLA; *SORL1*) downstream of the mitogen-activated protein kinase pathway. In addition, we demonstrate that SorLA interacts directly with HER3, forming a trimeric complex with HER2 and HER3 to attenuate lysosomal degradation of the dimer through a Rab4-dependent manner. In line with a role for SorLA in supporting the stability of the HER2 and HER3 receptors, loss of SorLA compromised heregulin-induced cell proliferation and sensitized metastatic anti-HER2 therapy-resistant breast cancer cells to neratinib in cancer spheroids *in vitro* and *in vivo* in a zebrafish brain xenograft model. Collectively, our results demonstrate a novel feed-forward loop consisting of heregulin, HER2-HER3 and SorLA, which controls breast cancer growth and anti-HER2 therapy resistance *in vitro* and *in vivo*.

**Significance:** HER3 signaling, through ERK/MAPK, upregulates SorLA and SorLA controls the trafficking and stability of HER3 to support cancer proliferation and neratinib resistance.

## Introduction

The human epidermal growth factor receptor (HER) family is composed of four transmembrane receptor tyrosine kinases (RTKs), encoded by the *EGFR* (HER1) and *ERBB2-4* (HER2-4) genes. These receptors signal through homo- and heterodimerization and promote cell transformation and oncogenic properties in multiple cancer types, including breast cancer (Junttila et al., 2009; Vaught et al., 2012; Yarden and Pines, 2012). Much of the focus in this area has been on EGFR and HER2, which are well-established tumor drivers and targets of effective anti-cancer therapeutics (Slamon et al., 2001; Yarden and Pines, 2012). In contrast, the role of HER3 is less understood, and has until recently been underappreciated. This is largely owing to the fact that HER3 has impaired kinase activity and its phosphorylation depends on dimerization with other RTKs (Junttila et al., 2009; Mishra et al., 2018). However, an increasing number of studies acknowledge HER3 as a key driver of carcinogenesis due to its unique ability to directly activate the phosphatidylinositol-3-OH kinase (PI3K)/protein kinase B (AKT) signaling pathway. Moreover, HER3 dimerization with HER2, which has the strongest kinase activity among all HER proteins, represents the most potent signaling receptor pair within the HER family (Amin et al., 2010; Garrett et al., 2011; Mishra et al., 2018; Sergina et al., 2007; Vaught et al., 2012).

HER3 drives resistance to targeted therapies in a wide range of solid tumors, including *ERBB2*-amplified (HER2-positive) breast cancer (Chandarlapaty et al., 2011; Kodack et al., 2017; Mishra et al., 2018). Increased HER3 expression compensates for HER2 tyrosine kinase inhibition, and HER3 activation by residual HER2 activity sustains oncogenic signaling (Amin et al., 2010; Garrett et al., 2011). In addition, HER3 growth factor ligands, heregulins (a.k.a. neuregulins), mediate resistance to the anti-HER2 monoclonal antibody trastuzumab and the dual HER2/EGFR tyrosine kinase inhibitor lapatinib (Gijsen et al., 2010; Xia et al., 2013). Therefore, better management of the disease would require efficient and safe targeting of the heregulin/HER2/HER3 signaling unit in tumors (Kang et al., 2014; Mishra et al., 2018). However, so far, none of the reported anti-HER3 therapy clinical trials have resulted in Food and Drug Administration (FDA) approval in any cancer type (Mishra et al., 2018).

Sortilin-related receptor (SorLA; *SORL1*) is a type-1 transmembrane sorting receptor that directs cargo proteins to spatially defined locations within the cell (Willnow et al., 2010). SorLA belongs to the family of vacuolar protein sorting 10 protein (VPS10P)-domain receptors (Willnow et al., 2008), and is well characterized for its protective role in Alzheimer’s disease, and for bolstering insulin signaling in adipose tissue, by regulating the traffic and biological function of the amyloid precursor protein and the insulin receptor, respectively (Andersen et al., 2005; Schmidt et al., 2016; Willnow et al., 2010). The SorLA carboxy-terminal tail harbors different sorting motifs, which bind to cytosolic adaptor proteins that regulate SorLA intracellular trafficking between the Golgi and the cell surface through endosomal compartments (Fjorback et al., 2012; Jacobsen et al., 2002). The role of intracellular trafficking in the spatiotemporal regulation of EGFR signaling is well established (Al-Akhrass et al., 2017; Caldieri et al., 2018). However, much less is known about the trafficking details influencing the oncogenic properties of HER2, or HER2-HER3 heterodimers (Bertelsen and Stang, 2014). We recently demonstrated that SorLA plays an important role in cancer, where it supports the oncogenic fitness of HER2 by orchestrating HER2 traffic to the plasma membrane, increasing signaling and proliferation in HER2-positive breast cancer (Pietilä et al., 2019).

This study aims to investigate the role of SorLA in mediating targeted therapy resistance in breast cancer, with a focus on the signaling by the HER2-HER3 oncogenic driver. We find that heregulins induce transcription of *SORL1 via* HER2-HER3 signaling to the mitogen-activated protein kinase (MAPK) pathway. In addition, we demonstrate that SorLA supports HER2-HER3 expression in a Ras-related protein Rab4-dependent manner. Furthermore, we demonstrate, for the first time, that this regulation involves a direct SorLA interaction with HER2-HER3 dimer. SorLA silencing inhibits 3D spheroid growth induced by heregulin-enriched stroma, and sensitizes metastatic breast cancer cells to the HER2/EGFR dual tyrosine kinase inhibitor neratinib in an *in vivo* xenograft model of brain tumors. This highlights SorLA as a potential target for the development of combination therapies aimed at overcoming HER3-mediated resistance of HER2-positive breast cancer patients to existing anti-HER2 therapies.

## Results

### Heregulins regulate SORL1 expression

We stratified 59 breast cancer cell lines based on *ERBB3* or *ERBB2* expression by mining the publicly available Cancer Cell Line Encyclopedia (CCLE) database (Barretina et al., 2012). Our statistical analyses indicated significantly higher SorLA mRNA (*SORL1*) expression in high compared to low *ERBB3*- and *ERBB2*-expressing cells (Figure 1A, Figure S1A). In addition, *SORL1* expression was higher in tumors exhibiting high *ERBB3* and *ERBB2* expression (Figure 1B; breast cancer patient data from the METABRIC study on the cBioportal database (Cerami et al., 2012; Gao et al., 2013; Pereira et al., 2016)). These findings suggested that HER2 and HER3 could positively regulate *SORL1* expression in breast tumors.

To assess this hypothesis, we explored the effect of HER2-HER3 signaling on *SORL1* expression by stimulating BT-474 cells with heregulin β-1 (Hrg β-1) over a 72 h time course. Hrg β-1 treatment triggered an increase in SorLA protein levels in a time-dependent manner (Figure 1C & D). As expected, it also activated AKT (AKT phosphorylation; pAKT) and the MAPK cascade (ERK phosphorylation; pERK) (Figure 1C) (Mishra et al., 2018). The increase in SorLA correlated with elevated *SORL1* mRNA levels in Hrg β-1-treated BT-474 and MDA-MB-361 cells (Figure 1E, Figure S1B-D). These findings indicate that ligand-induced HER3 signaling positively regulates SorLA expression on the transcriptional level. In addition to exogenous Hrg β-1 ligand stimulation, autocrine ligand secretion in BT-474 cells, stably overexpressing Hrg β-1 (Figure S1E), induced ERK and AKT phosphorylation (Figure 1F), *SORL1* mRNA (Figure S1F) and SorLA protein levels (Figure 1F & G).

**Fig. 1.**
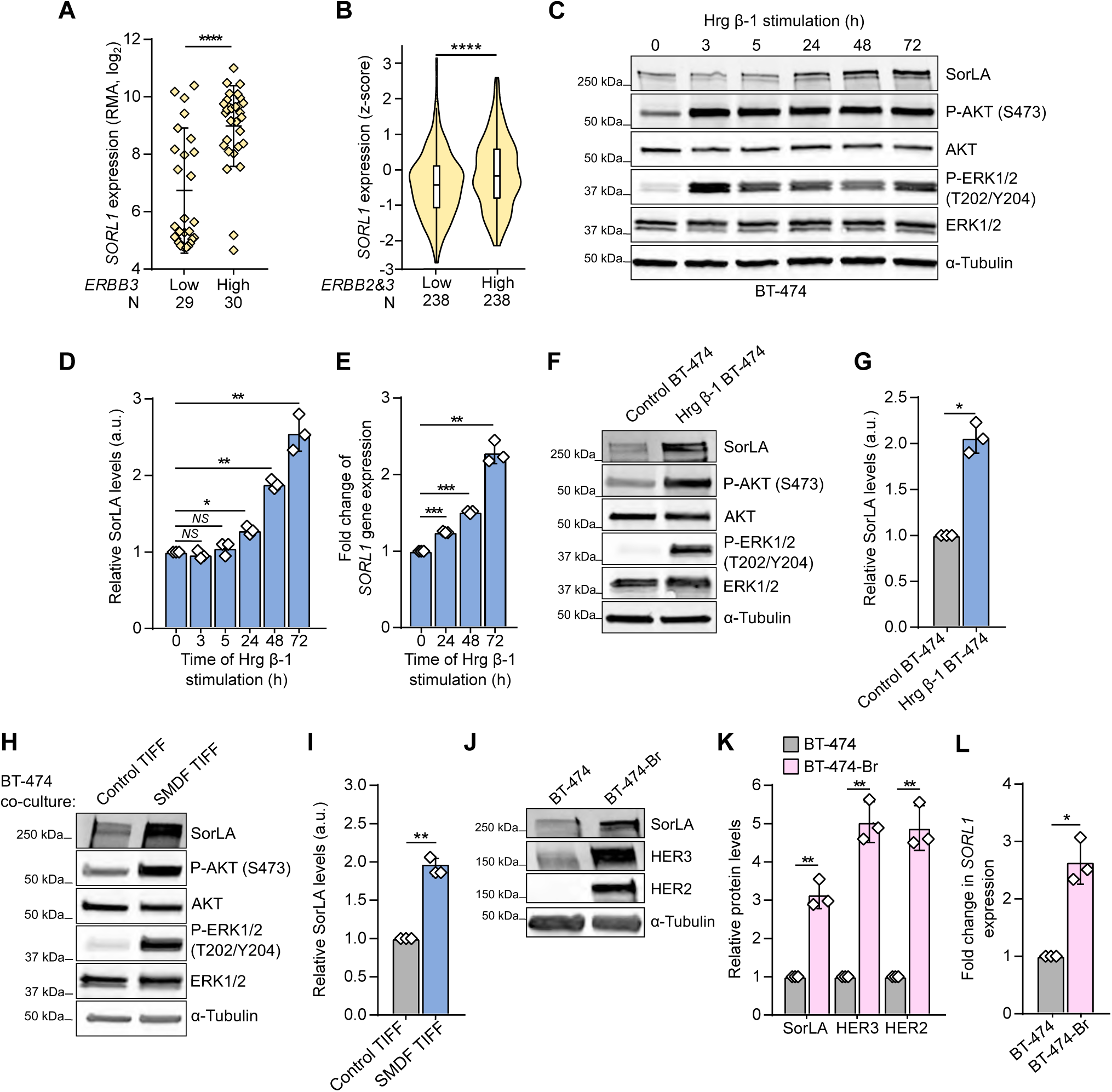
HER3 signaling regulates *SORL1* expression. **A**.*SORL1* expression is significantly higher in breast cancer cell lines with high *ERBB3* expression (CCLE; N=59). RMA: robust multi-array average. **B**. *SORL1* expression is higher in tumors with high *ERBB3* and *ERBB2* expression (cBioPortal; N=476). Violin plot boxes represent median and 25th and 75th percentiles (interquartile range), and whiskers extend to maximum and minimum values. **C**. Hrg β-1 increases SorLA levels. BT-474 cells were stimulated with 20 ng.mL^-1^ Hrg β-1 for the indicated times. Representative immunoblotting of SorLA, AKT(p)S473, total AKT, ERK1/2(p)T202/Y204, total ERK1/2, with αtubulin as a loading control. **D**. Quantification of SorLA levels normalized to loading control and relative to non-stimulated (0 h) cells. **E**. Hrg β-1 increases *SORL1* expression. Quantification of *SORL1* mRNA levels, relative to *HPRT1*, determined with RT-qPCR in BT-474 cells stimulated with 20 ng.mL^-1^ Hrg β-1 for the indicated time points relative to non-stimulated (0 h) cells. **F**. Representative immunoblotting of SorLA, AKT(p)S473, total AKT, ERK1/2(p)T202/Y204, total ERK1/2, with α-tubulin as a loading control from control (mCherry)- or Hrg β-1-overexpressing BT-474 cells. **G**. Quantification of SorLA levels normalized to loading control and relative to control cells. **H**. SMDF increases SorLA expression. BT-474 cells were cultured on a monolayer of mCherry-positive control or SMDF-overexpressing fibroblasts (TIFF). BT-474 cells were FACS sorted (see methods) and cell lysates were analyzed for SorLA, AKT(p)S473, total AKT, ERK1/2(p)T202/Y204, total ERK1/2, with α-tubulin as a loading control. **I**. Quantification of SorLA levels normalized to loading control and relative to control TIFF co-cultured cells. **J**. Representative immunoblotting of SorLA, HER2 and HER3, with α-tubulin as a loading control from parental BT-474 and brain-tropic metastasis variant BT-474-Br cells. **K**. Quantification of the indicated protein levels normalized to loading control and relative to BT-474 cells. **L**. Quantification of *SORL1* mRNA levels, normalized to *HPRT1*, determined with RT-qPCR in parental BT-474 and BT-474-Br cells relative to BT-474 cells. Data are mean ± SD from three independent biological experiments; statistical analysis: Student’s t-test (unpaired, two-tailed, unequal variance).

Hrg β-1 is an isoform resulting from alternative splicing of *NRG1* gene transcripts (Mei and Nave, 2014). To explore whether *SORL1* regulation is exclusive to Hrg β-1, we established a model of telomerase-immortalized foreskin fibroblasts (TIFF) with stable overexpression of SMDF (heregulin isoform 10), which exhibits neuronal functions (Mei and Nave, 2014) (Figure S1G). We found that co-culturing BT-474 cells with SMDF-TIFF significantly elevates SorLA levels and triggers AKT and ERK signaling in BT-474 cells (Figure 1H & I). In addition, exposing BT-474 cells to conditioned medium from SMDF-TIFF significantly induced *SORL1* levels (Figure S1H). This indicates that *SORL1* upregulation by HER3 signaling occurs both in a paracrine and in an autocrine manner, and is not restricted to a specific heregulin isoform.

To further validate the role of HER2-HER3 in augmenting *SORL1* expression, we used an *in vivo* established model of brain-trophic metastatic BT-474 cells (Zhang et al., 2013). BT-474 cells from brain metastases (BT-474-Br) expressed significantly higher HER3 and HER2 levels, compared to the parental BT-474 cells, and this correlated with increased SorLA protein expression as well as *SORL1* transcription (Figure 1J-L). Taken together, these results demonstrate that activation of HER2-HER3 positively regulate SorLA/*SORL1* expression in breast cancer.

### HER3 signaling to ERK1/2 upregulates SORL1 expression

Next, we investigated the mechanistic details of heregulin-induced *SORL1* upregulation. We generated reporter constructs by placing a series of *SORL1* proximal promoter sequences in front of the firefly luciferase (Figure 2A; P1-7). The P1-7 constructs were expressed individually in BT-474 cells, stimulated or not with Hrg β-1 for 24 h. Readouts of luciferase activity indicated that P3 is the minimum promoter sequence required for transcription in basal cell culture conditions (Figure 2B). In addition, P3 exhibited the highest increase in luciferase activity upon Hrg β-1 stimulation (Figure 2B), highlighting this region to contain responsive elements to Hrg β-1 stimulation. To identify the intracellular signaling pathway responsible for Hrg β-1-mediated *SORL1* expression, we tested the ability of different signaling inhibitors to reduce Hrg β-1-induced P3 luciferase activity. The inhibitors were selected to specifically target individual proteins within the PI3K/AKT/mammalian target of rapamycin (mTOR) and ERK pathways known to be activated downstream of HER3 upon ligand stimulation (Mishra et al., 2018). Trametinib, an ERK pathway inhibitor, significantly decreased Hrg β-1-induced luciferase activity of P3 (a similar trend was observed also with ERK kinase and ERK1/2 inhibitors selumetinib and SCH772984, respectively) (Figure S2). In contrast, PI3K, AKT and mTOR inhibitors significantly enhanced Hrg β-1-induced P3 activity (Figure S2). This may be linked to the notion that AKT and mTOR inhibition can trigger compensatory signaling through the ERK pathway (Carracedo et al., 2008; Mendoza et al., 2011) and indicates that ERK signaling positively regulates the promoter activity of *SORL1* in response to Hrg β-1. In line with these luciferase promoter activity data, trametinib decreased GFP intensity in Hrg β-1-stimulated cells that express GFP under the control of the P3 promoter sequence (Figure 2C & D). In addition, Hrg β-1-induced upregulation of SorLA protein in BT-474 cells was sensitive to trametinib, but not the AKT inhibitor MK-2206 (Figure 2E & F), and trametinib inhibited *SORL1* mRNA expression in Hrg β-1-stimulated cells (Figure 2G). Taken together these data uncover a previously unknown mecha-nism by which HER2-HER3 signaling to ERK1/2 regulates *SORL1* transcription leading to increased SorLA protein levels in breast cancer.

**Fig. 2.**
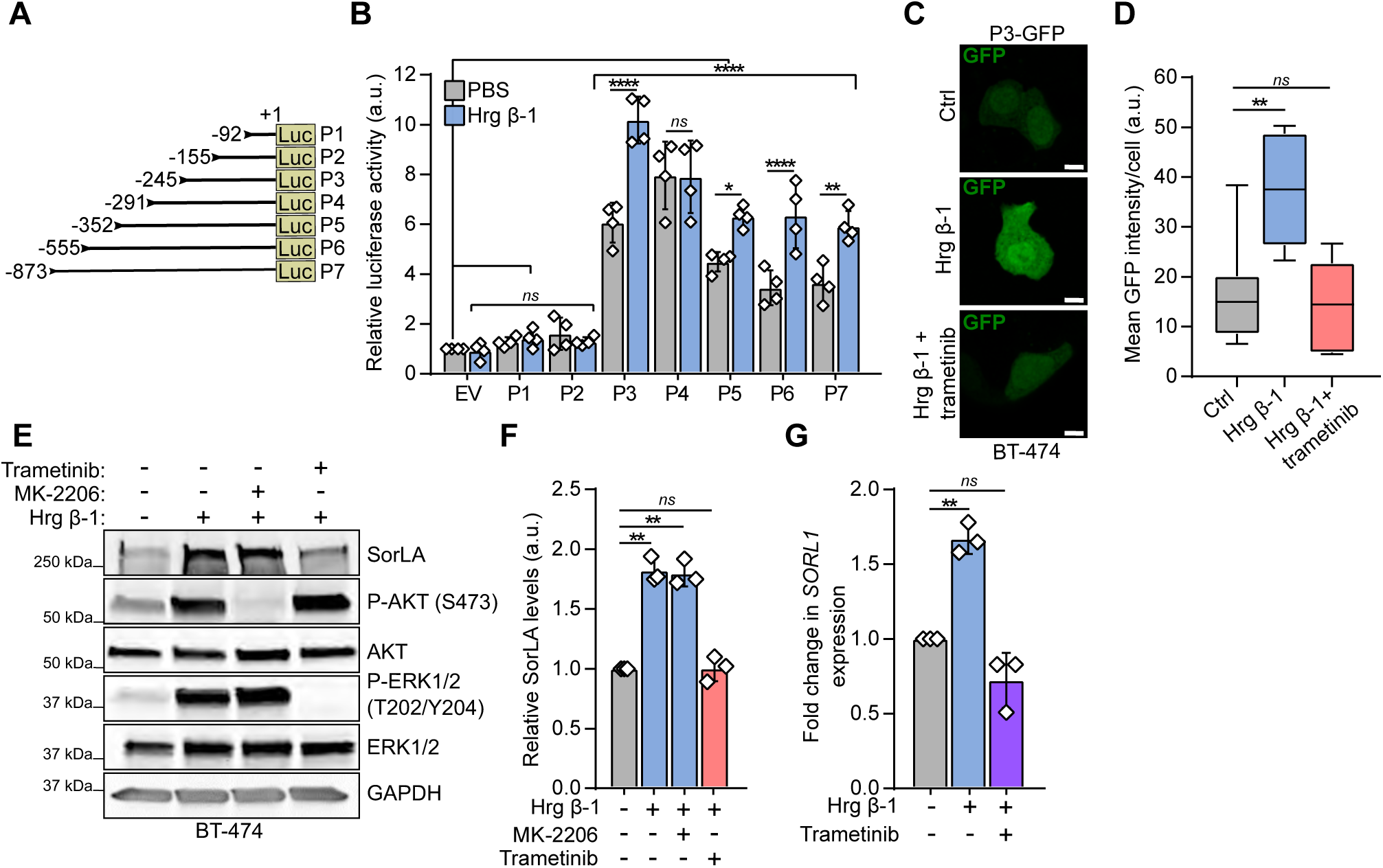
Heregulin-induced *SORL1* regulation requires HER3 signaling through ERK1/2. **A**. Representation of the different *SORL1* proximal promoter constructs (P1-7) used in luciferase assays with their corresponding molecular lengths (bp). **B**. P3 is a responsive promoter sequence to Hrg β-1 stimulation. A control empty vector (EV) and P1-7 were individually expressed in BT-474 cells together with pRL-TK Renilla luciferase transfection control and cells were stimulated or not with 20 ng.mL^-1^ Hrg β-1 for 24 h. Shown are relative luciferase activities to the control sample (EV, PBS-treated). **C**. Representative confocal microscopy images of P3-GFP promoter reporter construct-expressing BT-474 cells treated or not for 24 h with 20 ng.mL^-1^ Hrg β-1 in the presence or absence of 100 nM trametinib. Scale bars: 10 µm. **D**. Quantification of the mean intensity of the GFP signal per cell (whole-cell area). N=36 cells per group. Three biological replicates. **E**. Trametinib inhibits Hrg β-1-induced upregulation of SorLA. BT-474 cells were co-treated with 20 ng.mL^-1^ Hrg β-1 and either the pan-AKT inhibitor MK-2206 (2 µM) or the ERK pathway inhibitor trametinib (100 nM) for 48 h. Representative immunoblotting of SorLA, AKT(p)S473, total AKT, ERK1/2(p)T202/Y204, total ERK1/2, with GAPDH as a loading control. **F**. Quantification of SorLA levels normalized to loading control and relative to non-treated cells. **G**. *SORL1* mRNA levels, relative to *HPRT1*, determined with RT-qPCR in BT-474 cells stimulated or not with 20 ng.mL^-1^ Hrg β-1 in the presence or absence of trametinib relative to non-treated cells. B, F, G: data are mean ± SD from four (B) and three (F & G) independent biological replicates; statistical analysis: (B) Two-way ANOVA, Dunnett’s multiple comparisons test. (F & G) Student’s t-test (unpaired, two-tailed, unequal variance). (D) Box plots represent median and interquartile range, and whiskers extend to maximum and minimum values; One-way ANOVA, Dunn’s multiple comparisons test.

### SorLA regulates HER3 stability

SorLA regulates HER2 stability in breast cancer (Pietilä et al., 2019). Whether this regulation is exclusive to HER2 remained unknown. This prompted us to investigate the role of SorLA in regulating HER3 expression. We focused on HER3 since HER2-HER3 dimers control SorLA/*SORL1* expression (Figures 1 & 2) and drive therapy resistance in breast cancer (Amin et al., 2010; Mishra et al., 2018). As a first exploratory analysis, we assessed whether SorLA and HER3 protein levels correlate using the quantitative proteomics of the CCLE on the DepMap portal database (Nusinow et al., 2020). We found that SorLA and HER3 levels positively correlate across 29 breast cancer cell lines (Figure 3A). Next, we validated this correlation using a panel of six breast cancer cell lines exhibiting high, moderate and low SorLA levels (Figure 3B; Figure S3A).

**Fig. 3.**
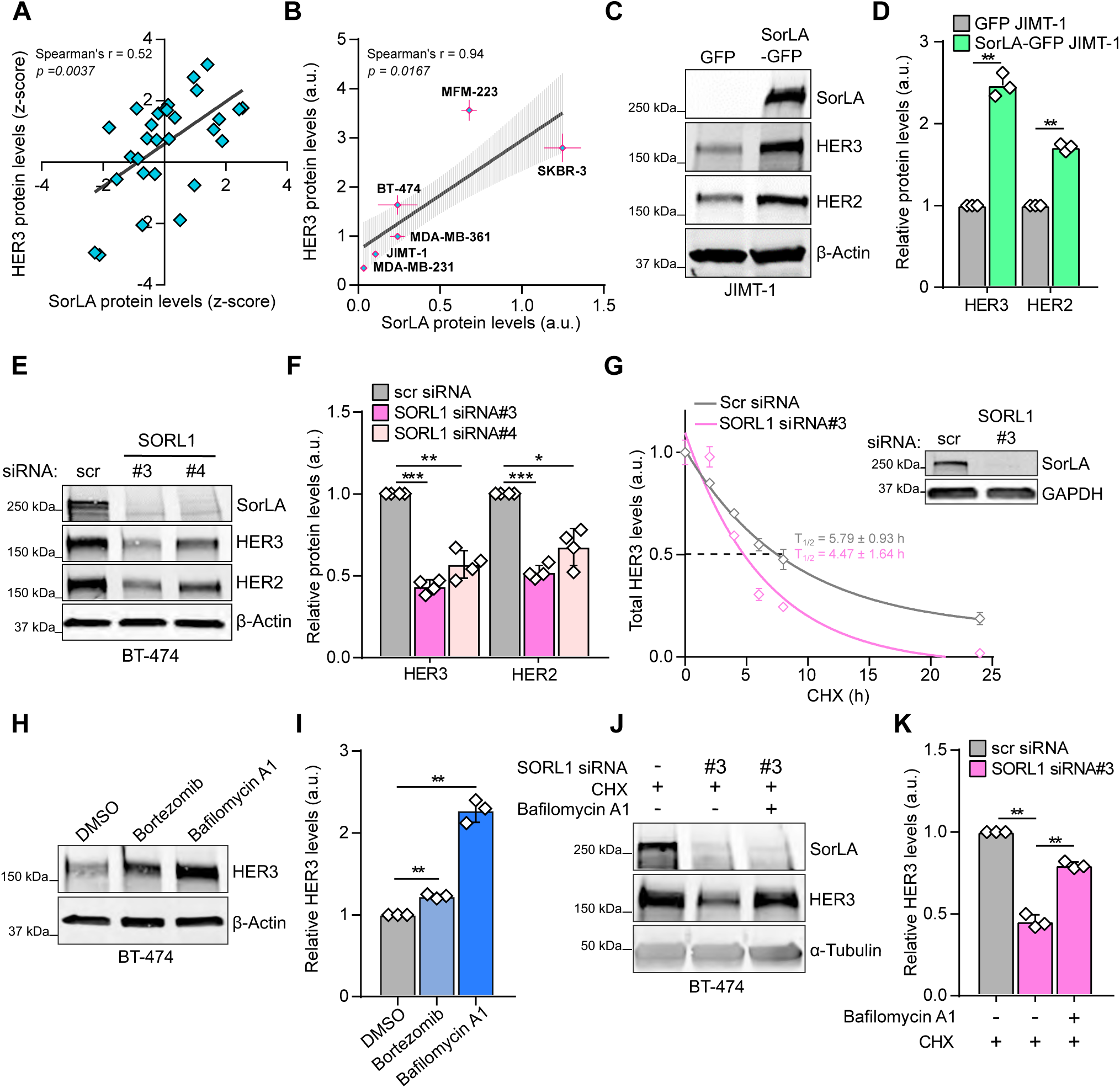
SorLA regulates HER3 stability. **A**. SorLA and HER3 protein levels correlate positively in breast cancer cell lines (DepMap portal; N=29). **B**. Correlation analysis between SorLA and HER3 protein levels in a panel of breast cancer cells. SorLA (x-axis) and HER3 (y-axis) levels (determined by Western blot, see Figure S3A) are expressed as the ratio of protein/α-tubulin control. r: correlation coefficient determined with non-parametric two-tailed Spearman test. **C-F**. HER3 and HER2 expression correlates with SorLA in HER2-positive breast cancer. **C & D**. SorLA-GFP transfection in JIMT-1 cells increases HER2 and HER3 levels compared to GFP transfected cells. **E & F**. SorLA silencing in BT-474 cells decreases HER2 and HER3. **C & E**. Representative immunoblotting of total HER2, HER3 and SorLA with β-actin as a loading control. **D & F** represent the respective quantifications of immunoblots in C & E with HER2/HER3 levels normalized to loading control and relative to control-silenced cells. **G**. SorLA silencing decreases HER3 stability. RNAi transfected BT-474 cells were treated with 25 µg.mL^-1^ of CHX for the indicated time points and HER3 protein levels were determined by immunoblotting (see Figure S3E). Shown are HER3 levels normalized to α-tubulin and relative to 0 h timepoint. Half-lives (T_1/2_) represent the time required for HER3 to decrease to 50% of its initial level. The least squares fitting method and extra-sum-of-squares F test were used to assess the statistical difference between curves from control and SorLA-silenced cells (*P=0*.*0002*). A representative western blot validating SorLA silencing is shown. **H**. HER3 is primarily degraded through the lysosomal pathway. BT-474 cells were treated with 1 µM of bortezomib or 50 nM of bafilomycin A1 for 4 h to inhibit proteasome and lysosome activities, respectively. HER3 expression was analyzed by immunoblotting, with α-tubulin as a loading control. **I**. Quantification of HER3 levels normalized to loading control and relative to DMSO-treated control cells. **J**. SorLA silencing triggers HER3 lysosomal degradation. SorLA-silenced BT-474 cells were co-treated for 4 h with CHX and bafilomycin A1. HER3 expression was analyzed by immunoblotting, with α-tubulin as a loading control. **K**. Quantification of HER3 levels normalized to loading control and relative to CHX-treated control-silenced cells. Data are mean ± SD from three (B, D, G, I, K) or four (F) independent biological replicates. Statistical analyses: Student’s t-test (unpaired, two-tailed, unequal variance) unless indicated otherwise. Scr: control non-targeting siRNA.)

To assess whether SorLA regulates HER3 in HER2-positive breast cancer, we expressed SorLA in JIMT-1 cells, which have very low endogenous SorLA expression compared to other HER2-positive cell lines (Pietilä et al., 2019). Expression of SorLA increased HER3 levels significantly (Figure 3C & D). Conversely, SorLA silencing in two endogenous SorLA-expressing HER2-positive cell lines, BT-474 and MDA-MB-361, with two different siRNAs significantly decreased HER3 expression (Figure 3E & F; Figure S3B & C). This indicates a regulation of HER3 by SorLA, in line with comparable effects of SorLA overexpression or silencing on HER2 levels (Figure 3C-E; Figure S3B & C), consistent with our previous findings (Pietilä et al., 2019).

To characterize the regulation of HER3 by SorLA in more detail, we performed qPCR analyses using our gain- and loss-of-function models. The expression of *ERBB3* remained unchanged upon SorLA overexpression or silencing, indicating that SorLA regulates HER3 at a posttranscriptional level (Figure S3D). These results prompted us to test whether SorLA regulates HER3 stability. Cycloheximide (CHX) chase experiments revealed a significantly shorter HER3 half-life (T_1/2_) in SorLA-silenced cells (T_1/2_ = 4.5 ± 1.6h) (Figure 3G; Figure S3E) compared to control-silenced cells (T_1/2_ = 5.8 ± 0.9h). Bortezomib-mediated inhibition of proteasome activity resulted in only a slight increase in HER3 levels, whereas, bafilomycin A1-mediated inhibition of lysosome function more than doubled HER3 protein levels (Figure 3H & I), indicating that HER3 primarily undergoes lysosomal degradation in BT-474 cells. Moreover, the enhanced HER3 degradation, observed in CHX-treated SorLA-silenced cells, could be largely rescued by bafilomycin A1 treatment (Figure 3J & K). Cumulatively, these data implicate SorLA in attenuation of HER3 lysosomal degradation.

### SorLA interacts with the HER2-HER3 dimer

Next, we aimed to explore whether the regulation of HER2 and HER3 stability is linked to SorLA association with the dimer. Bi-molecular fluorescence complementation (BiFC) is a method to detect protein-protein interactions in live cells and is based on the reconstitution of a fluorescent protein (Venus in this study) *via* reassembly of two truncated, and non-fluorescent, N- (v1) and C-terminal (v2) fragments, fused to interacting proteins of interest (Hu et al., 2002) (Figure 4A). Imaging of HER2-positive BT-474 cells co-expressing SorLA-v1 with either HER3-v2 or HER2-v2 revealed a strong, predominately intracellular fluorescent signal (Figure 4B, insets 1-4), indicating the formation of SorLA-HER2 and SorLA-HER3 complexes in cells. HER2-HER3 (fused to v2 and v1, respectively) complexes were detected both at the cell surface and inside the cells (Figure 4B, insets 5 & 6), demonstrating that HER2-HER3 heterodimers localize both to intracellular endomembrane-like structures and to the plasma membrane. To assess whether SorLA interacts with HER2-HER3 heterodimers, we used a conformation-specific nanobody to detect complemented Venus and to affinity purify the BiFC SorLA-HER3 complexes (Croucher et al., 2016). This biochemical approach revealed HER2 association with SorLA-HER3, indicative of the formation of a SorLA-HER2-HER3 trimeric complex in breast cancer cells (Figure 4C).

**Fig. 4.**
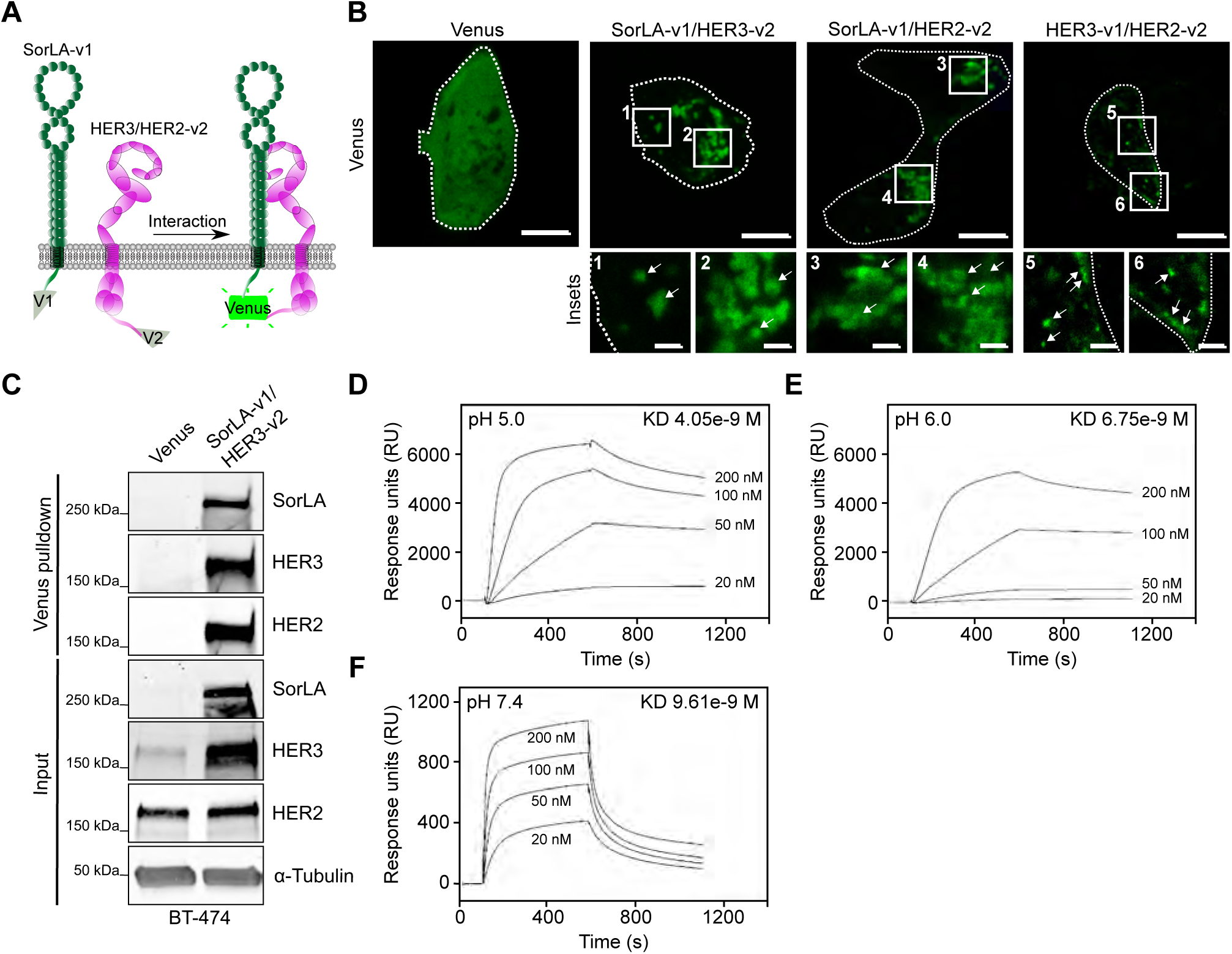
SorLA interacts with HER2-HER3 dimers. **A**. A scheme depicting BiFC between SorLA-v1 and HER2/3-v2. **B**. The indicated BiFC dimers were expressed in BT-474 cells and their interaction was assessed through imaging of reconstituted Venus. Cells are outlined with white dotted lines. Shown are representative confocal microscopy images and a control Venus-expressing cell showing diffuse fluorescence localization. Scale bars (main): 10 µm. Scale bars (insets): 2 µm. **C**. Venus or SorLA-v1/HER3-v2 were transiently expressed in BT-474 cells. Cell lysates were subjected to nanobody pulldown (specific for the reconstituted Venus v1+v2 dimer) and pulldowns and total cell lysates (input) were immunoblotted with the indicated antibodies. **D-F**. SorLA interacts with HER3 in a pH-dependent manner. SPR analysis on immobilized SorLA and a 20-200 nM concentration series of HER3 at pH 5.0 (D), 6.0 (E) and 7.4 (F). KD: equilibrium dissociation constant. Data are representative of three independent biological replicates.

To determine whether SorLA interacts directly with the extracellular domains of HER2 and HER3, we performed surface plasmon resonance (SPR) analysis, an assay that records protein interaction with an immobilized target on a microchip (Schuck, 1997). SPR analyses were performed using immobilized SorLA ectodomain under pH conditions corresponding to either endosomal (pH 5 & 6) or cell-surface compartments (7.4) (Maxfield and McGraw, 2004; Wang et al., 2015). Addition of the HER3 ectodomain at increasing concentrations triggered a rapid and reversible surge in binding (response units), indicating a specific interaction between SorLA and HER3 ectodomains (Figure 4D). The strength of this interaction decreased with increasing pH (Figure 4D-F), indicating that this interaction is pH-sensitive, similar to the interaction of SorLA with the amyloid precursor protein (Mehmedbasic et al., 2015). A similar interaction pattern was observed when increasing concentrations of the HER2 ectodomain were applied at different pH values (Figure S4A-C). The kinetics of these interactions are given as supporting information (Supplementary Table 1). The SPR results demonstrate a previously unappreciated direct interaction of SorLA with both HER2 and HER3, and enhanced interaction at a lower pH, characteristic of endosomes.

### SorLA regulates HER2 and HER3 stability in a Rab4-dependent fashion

SorLA regulates HER2 stability by supporting recycling of the receptor to the plasma membrane (Pietilä et al., 2019). However, the mechanistic details of this process and its implications for HER3 remained to be determined. To investigate the trafficking pathway underpinning this regulation, we first assessed intracellular co-localization between GFP-SorLA and different RFP-tagged endosomal markers: early endosome antigen-1 (EEA1); vacuolar protein sorting-associated protein 29 (VPS29), a subunit of the retromer complex, which mediates endosome-to-Golgi receptor retrieval (Small and Petsko, 2015); and Rab4 and Rab11, which regulate receptor recycling from early endosomes and the endocytic recycling compartment, respectively (Grant and Donaldson, 2009). Using confocal microscopy (Figure S5A) and super–resolution Airyscan confocal microscopy (Figure 5A), we readily detected overlapping signals between SorLA and the tested endosomal markers. Co-localization analysis indicated the highest degree of co-localisation between SorLA and Rab4 (Figure 5A, insets 3 & 4; Figure 5B; Figure S5A, insets 3 & 4) indicating that SorLA may function in Rab4-containing recycling endosomes in BT-474 cells. Indeed, overexpression of a dominant-negative GDP-locked Rab4^S22N^ inhibited SorLA expression-induced up-regulation of HER2 and HER3 in SorLA-low JIMT-1 cells (Figure 5C & D). This indicates that SorLA regulates HER2 and HER3 expression in a Rab4-dependent manner. In line with these results, we found that SorLA-, HER2- and HER3-containing BiFC dimers reside in Rab4-positive intracellular compartments in BT-474 cells (Figure 5E & F), further highlighting the Rab4 pathway in mediating SorLA regulation of HER2-HER3 complexes in breast cancer.

**Fig. 5.**
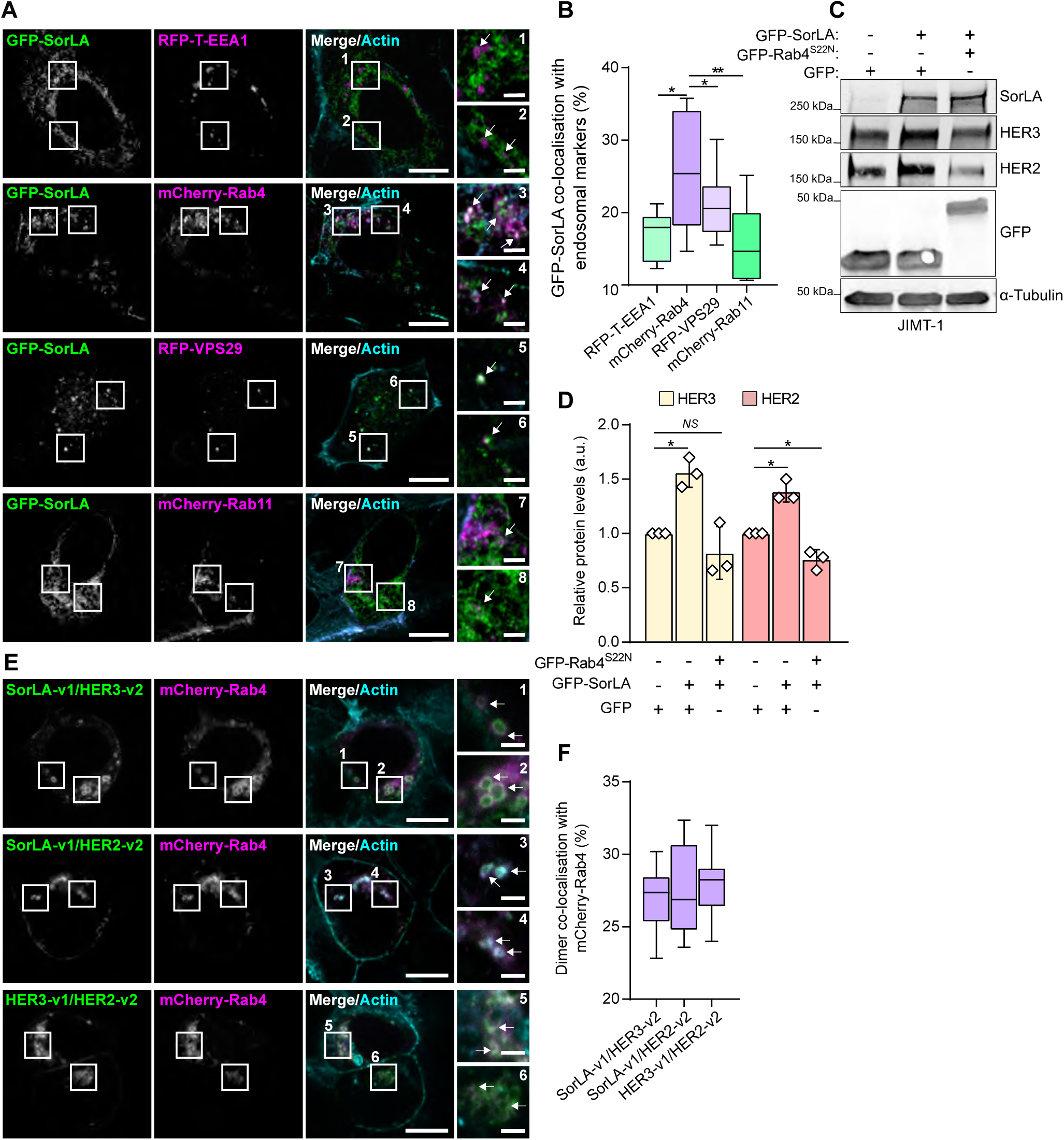
SorLA regulation of HER2 and HER3 requires functional Rab4. **A**. Representative Airyscan confocal microscopy images of BT-474 cells co-expressing GFP-SorLA with the indicated endosomal markers. SiR-Actin was used for counterstaining the actin cytoskeleton. White arrows depict co-localizing signals. Scale bars: 10 µm. Scale bars (insets): 2µm. **B**. GFP-SorLA strongly co-localizes with mCherry-Rab4 in BT-474 cells. Co-localization was calculated (see methods) from BT-474 cells transfected and imaged as in (A). N=36 cells per group. **C**. JIMT-1 cells were co-transfected with GFP-SorLA and either GFP control or GFP-Rab4S22N dominant-negative mutant. Representative immunoblotting of SorLA, HER2, HER3 and GFP, with α-tubulin as a loading control. **D**. Quantification of HER2 and HER3 levels normalized to loading control and relative to control GFP-transfected cells. **E**. Representative confocal microscopy images of BT-474 cells co-overexpressing mCherry-Rab4 with the indicated BiFC dimers. SiR-Actin was used for counterstaining the actin cytoskeleton. White arrows depict co-localizing signals. Scale bars: 10 µm. Scale bars (insets): 2 µm. **F**. Co-localization analysis between BiFC and mCherry-Rab4. N=30 cells per group. (D) Data are mean ± SD from three independent biological experiments; statistical analysis: Student’s t-test (unpaired, two-tailed, unequal variance). (B & F) Box plots represent median and 25th and 75th percentiles (interquartile range), and whiskers extend to maximum and minimum values; three biological replicates. Statistical analysis: One-way ANOVA, Dunn’s multiple comparisons test.

### SorLA is necessary for HER3-driven oncogenic cell growth

Our data thus far indicate a feed-forward loop between HER3 and SorLA where HER3 signaling induces SorLA expression and SorLA supports HER3 stability. Next, we evaluated the role of SorLA in the phenotype of HER3-driven cancer. Hrg β-1 stimulation resulted in enhanced cell viability in control-silenced BT-474 cells (Figure 6A), whereas SorLA silencing resulted in diminished cell viability irrespective of Hrg β-1 stimulation (Figure 6A). In addition, SorLA silencing, with two distinct siRNAs, inhibited the viability of Hrg β-1-expressing BT-474 cells (Figure 6B). These data indicate that SorLA is required for Hrg β-1-induced cell viability in 2D cell culture. Next, we investigated the role of SorLA in modulating heregulin effects within a more physiologically representative experimental setting. Control- and SorLA-silenced mCherry-BT-474 cells were co-cultured as spheroids with either control or Hrg β-1-overexpressing fibroblasts, using a matrigel-based 3D culture system. While co-culture with Hrg β-1-overexpressing TIFF strongly promoted the growth of control-silenced BT-474 spheroids, it had no effect on SorLA-silenced BT-474 spheroids (Figure 6 C & D). These results indicate that SorLA expression is essential for HER3-driven growth of tumor spheroids in heregulin-enriched stroma.

**Fig. 6.**
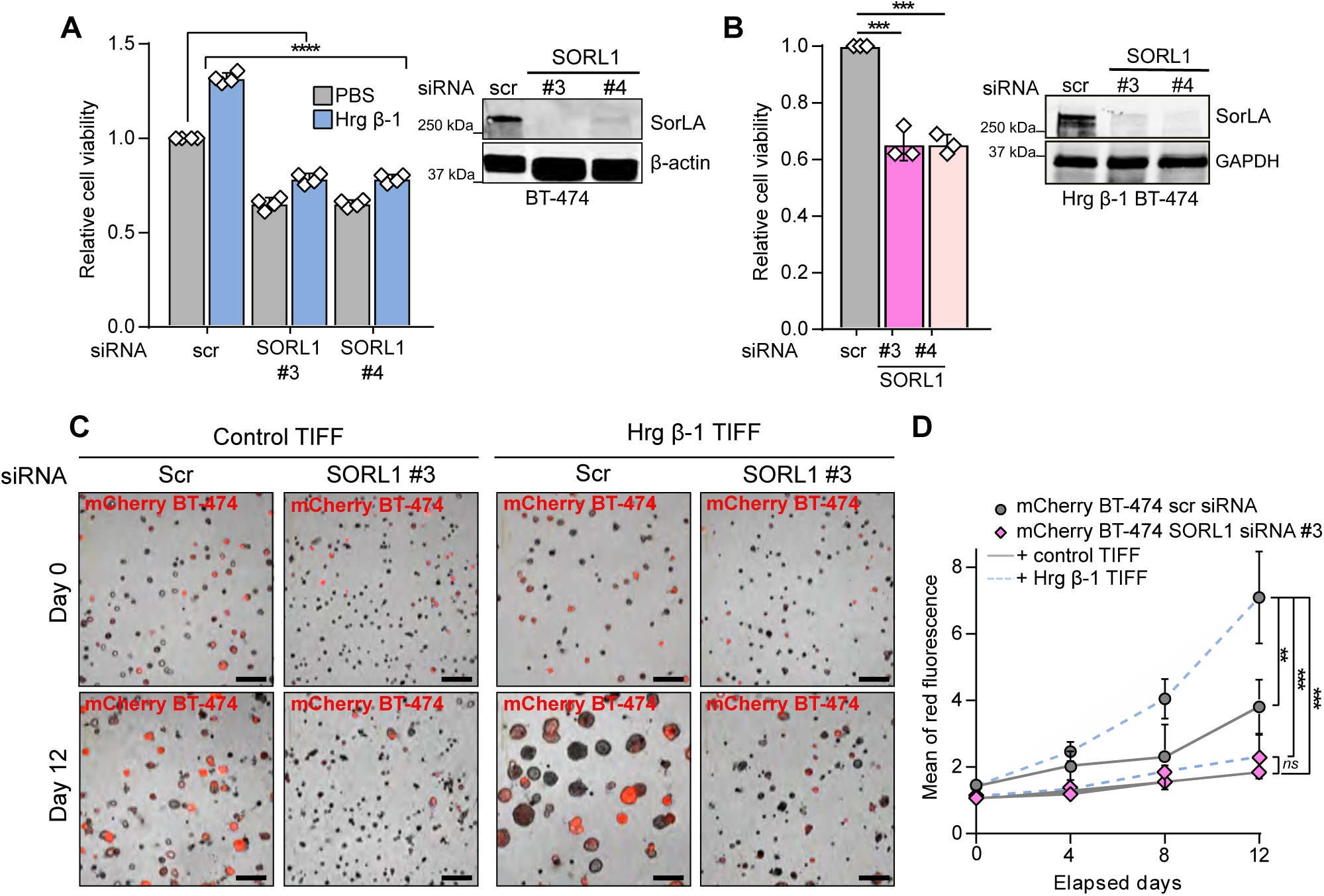
SorLA expression determines heregulin response. **A**. SorLA silencing decreases cell viability in basal and Hrg β-1-enriched cell culture conditions. Control and SorLA-silenced BT-474 cells were stimulated or not with 20 ng.mL^-1^ Hrg β-1 for 48 h and cell viability was assessed using a WST-8-based method. Cell viability values are represented as fold change relative to unstimulated control BT-474 cells. A representative western blot validating SorLA silencing is shown. **B**. Hrg β-1-expressing BT-474 cells were silenced for SorLA and cell viability was analyzed as in (A). A representative western blot validating SorLA silencing is shown. **C & D**. SorLA silencing inhibits spheroid growth in Hrg β-1-enriched extracellular matrix. SorLA-silenced mCherry BT-474 cells were co-cultured with Hrg β-1-overexpressing fibroblasts (Hrg β-1 TIFF) in matrigel for 12 days. Shown are representative live-cell images (C) and quantification of mCherry fluorescence reflecting spheroid growth (D). Data are mean ± SD from three independent biological replicates; statistical analysis: A & D, One-way ANOVA, Dunn’s multiple comparisons test and B, Student’s t-test (unpaired, two-tailed, unequal variance). Scr = control non-targeting siRNA.

### SorLA silencing sensitizes resistant cells to neratinib

Increased HER3 activation is implicated in resistance to targeted therapeutics against HER2, PI3K and AKT in breast cancer (Chandarlapaty et al., 2011; Kodack et al., 2017; Xia et al., 2013). Hence, we speculated that in such targeted therapeutic settings SorLA silencing might have a beneficial effect on drug response. We chose MDA-MB-361 cells as a model as they are derived from a HER2-targeted therapy-resistant brain metastasis (Ding et al., 2018) and their HER2 and HER3 levels are SorLA dependent (Figure S3B & C). MDA-MB-361 cells were treated with 500 nM of either the dual HER2/EGFR tyrosine kinase inhibitor neratinib, the pan-class I PI3K inhibitor buparlisib or the pan-AKT inhibitor MK-2206 for 48 h. Consistent with its ability to inhibit PIK3CA-Mutant p110α regulatory subunit of PI3K (Baselga et al., 2017), buparlisib inhibited the viability of MDA-MB-361 control cells (Figure 7A). In accordance with the partial sensitivity of MDA-MB-361 cells to AKT inhibitors (Liu et al., 2018), MK-2206 triggered a slight decrease in cell viability (Figure 7A). No significant effect on cell viability was observed upon neratinib treatment of control MDA-MB-361 cells (Figure 7A). SorLA silencing, with two different siRNAs, inhibited MDA-MB-361 cell viability in basal cell culture conditions and SorLA silencing and neratinib exhibited synergistic inhibitory effects on MDA-MB- 361 cell growth (Figure 7A). We did not observe synergistic effects using buparlisib and MK-2206 (Figure 7A) indicating that SorLA silencing specifically alters the response to neratinib.

**Fig. 7.**
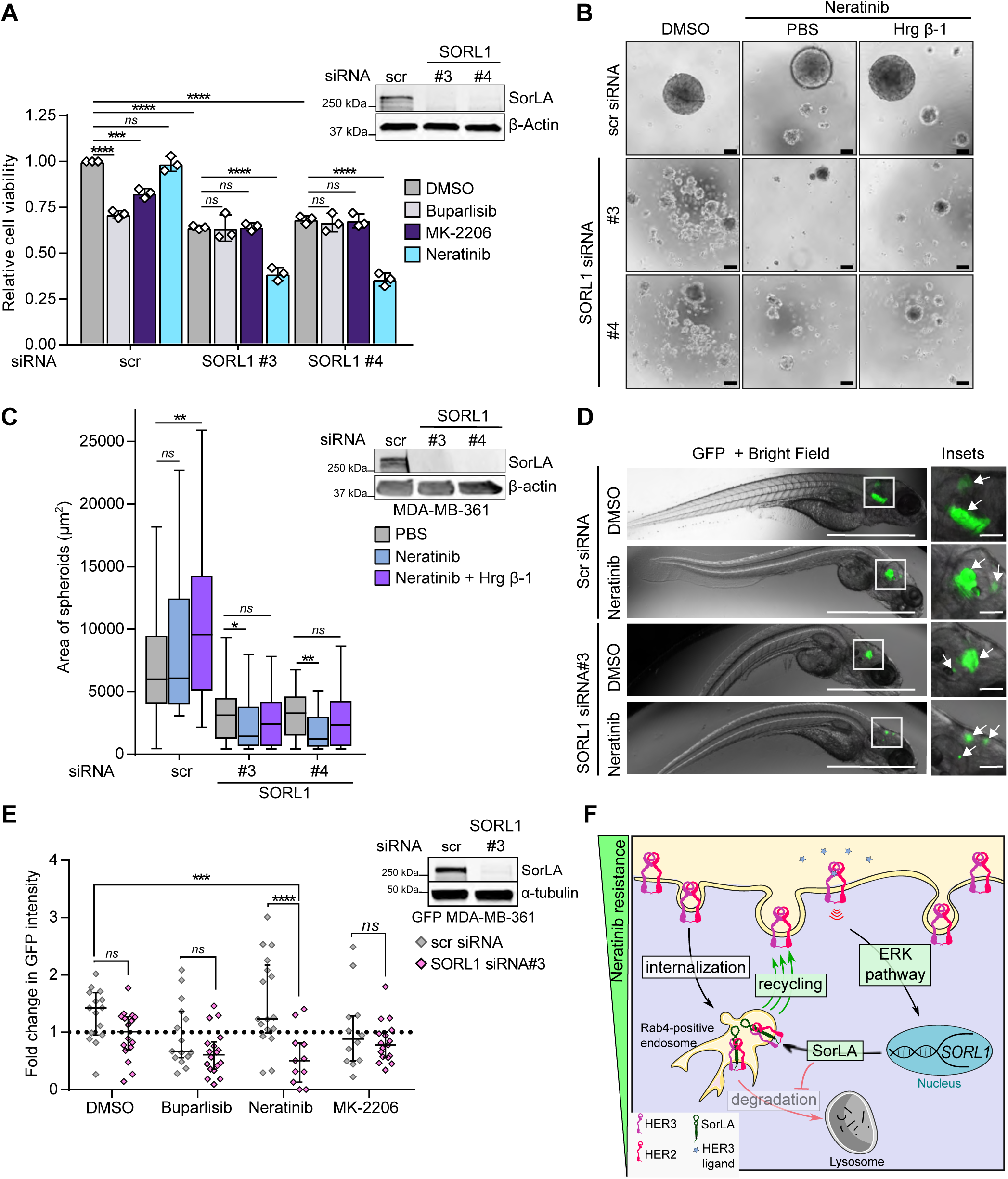
SorLA silencing specifically reverts resistance to neratinib *in vitro* and *in vivo*. **A**. SorLA silencing and neratinib show synergistic effects in inhibiting MDA-MB-361 cell growth. Control- and SorLA-silenced MDA-MB-361 cells were treated with 500 nM of either the pan-PI3K inhibitor buparlisib, the pan-AKT inhibitor MK-2206 or the dual HER2/EGFR tyrosine kinase inhibitor neratinib for 48 h. Cell viability was measured using WST-8-based method. Values are represented as fold change relative to DMSO-treated control MDA-MB-361 cells. A representative western blot validating SorLA silencing is shown. **B & C**. SorLA-silencing exhibits a synergistic effect with neratinib in inhibiting anchorage-independent spheroid growth of MDA-MB-361 cells in 3D low-attachment cell culture conditions. Control- and SorLA-silenced MDA-MB-361 cells were grown in low-attachment cell culture conditions for 7 days in the presence of the indicated treatments. Scale bar 100 µm. Spheroid sizes are quantified in and a representative western blot validating SorLA silencing are shown in (C). N>50 spheroids per group. **D**. Control and SorLA-silenced GFP-MDA-MB-361 cells were engrafted in the brain of zebrafish embryos and allowed to grow for 4 days in the presence of DMSO control or neratinib (400 nM). Representative GFP + Bright Field images of brain tumors are shown. Scale bars: 1 mm. Scale bars (insets): 100 µm. **E**. Control- and SorLA-silenced GFP-MDA-MB-361 cells were engrafted in zebrafish brain and allowed to grow for 4 days in the presence of DMSO, buparlisib (2 µM), neratinib (400 nM) or MK-2206 (400 nM). Tumor growth is represented as fold change in GFP intensity relative to day 1 post-engraftment. A representative western blot validating SorLA silencing is shown. **F** A representative scheme of a neratinib resistance mechanism driven by a feed-forward loop supporting SorLA-HER2-HER3 expression. Key elements of this loop are highlighted in green boxes. HER2-HER3 signaling increases *SORL1* expression through the ERK pathway. Increased SorLA levels prevent HER2-HER3 lysosomal degradation (red arrow with lined arrowhead) presumably by stimulating receptor recycling from Rab4-positive endosomes (green arrows). Data are (A), mean ± SD from three 3 independent biological replicates; (C), Box plots representing median and interquartile range; whiskers extend to maximum and minimum values, and (E), median with interquartile range. Statistical analysis: (A), two-way ANOVA, Dunnett’s multiple comparisons test. (C), One-way ANOVA, Dunn’s multiple comparisons test. (E), two-way ANOVA, Dunnett’s multiple comparisons test (main SorLA silencing effect *P*<*0*.*0001*). Scr = control non-targeting siRNA.

Since resistance to HER2 inhibition correlates with anchorage-independent growth (Boulbes et al., 2015), we analyzed the growth of SorLA-silenced MDA-MB-361 spheroids in neratinib-containing ultra-low attachment cell culture conditions. Control MDA-MB-361 spheroids were resistant to neratinib, and these neratinib-treated cells were able to grow in the presence of Hrg β-1 (Figure 7B & C). In contrast, SorLA silencing inhibited MDA-MB-361 spheroid growth, and the addition of neratinib further diminished the spheroid size even in spheroids treated with neratinib and Hrg β-1 (Figure 7B & C), demonstrating that SorLA silencing alters neratinib sensitivity in spheroids and that SorLA is essential for inducing heregulin effects on cell growth.

To determine whether loss of SorLA alters resistance to targeted therapy *in vivo*, we used a zebrafish model, an increasingly widely appreciated powerful tool in cancer research (Fazio et al., 2020). GFP-labeled MDA-MB-361 cells, transiently silenced for SorLA expression, were engrafted in the brain of zebrafish embryos and the fish were then treated with buparlisib, neratinib or MK-2206. Neratinib treatment of SorLA-silenced cells resulted in regressed tumor growth while control tumors remained neratinib-resistant (Figure 7D & E). In contrast, SorLA-silencing did not alter the response to buparlisib nor to MK-2206 (Figure 7E). This highlights SorLA silencing as a sensitizing approach for HER2-targeted therapy, established here in MDA-MB-361 cells, but providing an essential proof-of-principle for future efforts to target SorLA as part of novel combination therapies.

## Discussion

Here, we discovered a heregulin-dependent HER3 oncogenic signaling nexus, which forms the basis of a feed-forward loop supporting SorLA, HER2, and HER3 levels in breast cancer cells to drive neratinib resistance (Figure 7F). We identified that heregulin-mediated signaling activates *SORL1* transcription *via* ERK-dependent induction of the *SORL1* promoter. In addition, we unraveled mechanistic details of HER3 regulation by SorLA. We found that SorLA interacts directly, and in a pH-sensitive manner, with the HER2-HER3 heterodimer to support receptor stability at the protein level. We detected this interaction in Rab4-positive endosomes, which appear to be crucial intracellular compartments for SorLA to divert HER2-HER3 from lysosomal degradation. Thus, we have uncovered a positive feedback mechanism whereby increased SorLA levels support HER2-HER3 dimer signaling to drive cell proliferation, anti-HER2 therapy resistance and further increase SorLA expression in cells.

Heregulins are a family of growth factors encoded by 6 individual genes (*NRG1-6*) with *NRG1* representing the archetypical growth factor ligand associated with poor prognosis in HER2-positive breast cancer (Falls, 2003; Mei and Nave, 2014; Xia et al., 2013). The brain microenvironment is highly enriched with heregulins (Kodack et al., 2017). We found that the brain-trophic variants of BT-474 cells (Zhang et al., 2013) exhibit increased SorLA/*SORL1* expression. This raises the possibility that SorLA may be relevant in breast cancer brain metastases. Heregulin affects cell proliferation in a cell-type specific manner. Hrg β-1 increases the proliferation of BT-474 cells at relatively low concentrations, while it exhibits a suppressive growth effect on MDA-MB-361 cells (Hutcheson et al., 2011). Despite this, the two cell lines showed a similar increase in SorLA/*SORL1* expression upon heregulin stimulation, and both autocrine and paracrine signals by various heregulin proteins increased SorLA/*SORL1* expression. This suggests that the regulation of SorLA, downstream of HER2-HER3 in response to heregulin-enriched tissue, is a general regulatory mechanism in breast cancer.

Our data indicate a Rab4-dependency of SorLA-mediated stabilization of HER2 and HER3. This would be in line with the role of Rab4 in mediating recycling of EGFR (Tomas et al., 2014) and a recent study characterizing the role of altered Rab4-positive endosomes in sustaining EGFR signaling (Tubbesing et al., 2020). Nevertheless, the trafficking machinery linking SorLA to Rab4 remains to be investigated. The SorLA carboxy-terminal tail interacts with multiple trafficking proteins including GGA1 and GGA2 (Golgi-localizing, γ-adaptin ear homology domain, ARF-interacting proteins) (Jacobsen et al., 2002). GGA3 mediates Met RTK recycling from Rab4-positive endosomes (Parachoniak et al., 2011) suggesting the possibility that members of GGA family might influence SorLA-regulated RTK trafficking in HER2-positive breast cancer cells. In addition, defining the interactome of SorLA in complex with RTKs, in an unbiased manner, might uncover key novel adaptors/facilitators of SorLA-regulated traffic of receptors in cancer.

Our observation that heregulins upregulate SorLA and that SorLA, in turn, determines the heregulin response in BT-474 cells, alludes to a potential SorLA-dependent mechanism enabling metastatic breast cancer cells to adapt to, and colonize, the brain microenvironment. The heregulin-enriched brain parenchyma is known to promote resistance to anti-HER2 therapies enhancing the incidence of brain metastases that occurs in 50% of HER2-positive breast cancer patients (Kodack et al., 2012, 2015; Olson et al., 2013). Therefore, a molecular-level understanding of HER2-HER3 regulation in cells is required to not only dissect mechanisms of therapy relapse but to also provide alternative therapeutic options at such an advanced stage of the disease (Amin et al., 2010; Kang et al., 2014; Kodack et al., 2017; Mishra et al., 2018). A future therapeutic strategy undertaking an unbiased screening approach to identify potent SorLA blocking antibodies might provide a way forward in targeting heregulin-driven activation of HER2-HER3 dimer in breast cancer. Our study demonstrates that SorLA silencing alters resistance of HER2-positive breast cancer cells to neratinib in the zebrafish heregulin-enriched brain microenvironment (Sato et al., 2015). Neratinib is a dual HER2/EGFR tyrosine kinase inhibitor that was recently approved by the FDA for treatment of advanced or metastatic HER2-positive breast cancer (Research, 2020). We demonstrate that SorLA silencing exhibits a synergistic effect to neratinib, but not to buparlisib, or MK-2206. This might be linked to the ability of neratinib to induce ubiquitination and subsequent lysosomal degradation of HER2 (Zhang et al., 2016). Since SorLA silencing triggers HER2-HER3 lysosomal degradation, neratinib treatment could potentiate oncogenic-receptor targeting to the lysosomal pathway, a mechanism that does not apply to the other tested targeted chemotherapy agents. Given that HER3 drives therapy resistance and, despite extensive efforts, no anti-HER3 therapy is yet approved by the FDA (Mishra et al., 2018), targeting key regulators of HER3 stability, such as SorLA, might reveal a new field of research for drug discovery.

In summary, we describe an original role for SorLA as a positive regulator of the functional oncogenic driver HER2-HER3 in breast cancer. Since SorLA expression was associated with maintenance of anti-HER2 therapy resistance in brain metastasis xenografts, it may be a potential target for combating drug resistance. Additional research assessing the druggability of SorLA in breast cancer is warranted based on these findings.

## Materials and methods

### Cell culture and reagents

BT-474 (ATCC, HTB-20) and brain seeking BT-474-Br (generously provided by Dihua Yu (MD Anderson Cancer Center)) cells were grown in RPMI-1640 (Sigma-Aldrich, R5886) supplemented with 10% fetal bovine serum (FBS; Sigma-Aldrich, F7524), 1% vol/vol penicillin/streptomycin (Sigma-Aldrich, P0781-100ML) and L-glutamine. MDA-MB-361 cells were grown in Dulbecco’s modified essential medium (DMEM; Sigma-Aldrich, D5769) supplemented with 20% FBS, 1% vol/vol penicillin/streptomycin and L-glutamine. Telomerase immortalized foreskin fibroblasts (TIFF, generously provided by J. Norman (Beatson Institute for Cancer Research)) and JIMT-1 (DSMZ, ACC 589) cells were grown in DMEM supplemented with 10% FBS, 1% penicillin/streptomycin and L-glutamine. All cells were cultured in a humidified incubator set at 5% CO2 and 37°C. All cells were tested bimonthly – every 2 months – to ensure mycoplasma-free cell culture using MycoAlert™ mycoplasma detection kit (Lonza, #LT07-418) and MycoAlert™ assay control set (#LT07-518). The antibodies used are described in Supplementary Table 2. Previously published plasmids used in this study are summarized in Supplementary Table 3.

### Western blot

Cells were washed with ice-cold Dulbecco’s phosphate-buffered saline (DPBS, Gibco™, 11590476) prior to lysis with cell lysis buffer (CST, #9803) supplemented with 1% protease/phosphatase inhibitor cocktail (CST, #5872). Cell lysates were sonicated and cleared by centrifugation at 18,000×g for 10 min. Unless otherwise indicated, 30 µg of cleared lysates were subjected to SDS-PAGE under denaturing conditions (4–20% Mini-PROTEAN TGX Gels) and were transferred to nitrocellulose membranes (Bio-Rad Laboratories). Membranes were blocked with 5% milk-TBST (Tris-buffered saline and 0.1% Tween 20) and incubated with the indicated primary antibodies overnight at +4°C. Primary antibodies were diluted in blocking buffer (Thermo, StartingBlock (PBS) blocking, #37538) and PBS (1:1 ratio) mix and incubated overnight at +4 °C. After primary antibody incubation, membranes were washed three times with TBST and incubated with fluorophore-conjugated secondary antibodies diluted (1:2000) in blocking buffer at room temperature for 1h. Membranes were scanned using an infrared imaging system (Odyssey; LI-COR Biosciences). The following secondary antibodies were used: donkey anti-mouse IRDye 800CW (LI-COR, 926-32212), donkey anti-mouse IRDye 680RD (LI-COR, 926-68072), donkey anti-rabbit IRDye 800CW (LI-COR, 926-32213) and donkey anti-rabbit IRDye 680RD (LI-COR, 926-68073). The band intensity of each target was quantified using ImageJ (NIH) (Schneider et al., 2012) and normalized to loading control band intensity in each lane.

### Venus pull-down

Cells were lysed in Pierce IP Lysis Buffer (Thermo Scientific, 87787) supplemented with 1% protease/phosphatase inhibitor cocktail (CST, #5872). Lysates were cleared by centrifugation at 18,000×*g* for 10 min. 5% of cleared lysates were used as input control. Cleared lysates were incubated with GFP (Venus)-trap beads (Chromotek, gtak-20) for 50 min at +4°C to pull-down Venus-tagged proteins. Venus-trap beads were then pelleted by centrifugation at 3000 rpm for 3 min and washed 3 times with an IP wash buffer (20 mM tris-HCl (pH 7.5), 150 mM NaCl, 1% NP-40). Finally, pellets were boiled at 95°C for 10 min in sample buffer prior to SDS-PAGE.

### Quantitative reverse transcription-PCR

Total RNA was extracted according to the manufacturer’s instructions (NucleoSpin RNA extraction kit, MachereyNagel, 740,955.5). RNA concentration was measured by NanoDrop Lite (Thermo). Single-stranded cDNA was prepared using the high capacity cDNA reverse transcription kit (Applied Biosystems). The reaction was stopped by incubation at 95°C for 5 min. Approximately 100 ng of cDNA was used for each PCR reaction performed with TaqMan probes according to the manufacturer’s instructions (Thermo/Applied Biosystems, Taq-Man™ Universal Master Mix II, 4440040). The following TaqMan probes were used: SORL1 (Hs00268342_m1), ERBB3 (Hs00176538_m1), NRG1 (Hs00247624_m1) and HPRT1 (Hs02800695_m1). NRG1 primers recognize both Hrg β-1 and SMDF splicing variants of NRG1 gene transcripts. Relative quantification of gene expression values was calculated using the Δ Δ Ct method (Heid et al., 1996).

### Transient transfections

For transient overexpression, cells were transfected 24 h before experiments using Lipofectamine 3000 (Invitrogen, P/N 100022052) and P3000 enhancer reagent (Invitrogen, P/N100022058) according to the manufacturer’s instructions. For interference assays, cells were transfected 72 h before experiments using Lipofectamine RNAiMAX reagent (Invitrogen, P/N 56532) according to the manufacturer’s instructions. SORL1-targeting siRNAs were obtained from Dharmacon – siSORL1 #3 (J-004722-07, (5’CCGAA-GAGCUUGACUACUU3’)), siSORLA #4 (J-004722-05, (5’CCACGUGUCUGCCCAAUUA3’)). Allstars (Qiagen, 1027281) was used as a negative control. All siRNAs were used at a final concentration of 20 nM.

### Cell viability assays

Cells were silenced for SorLA in 6-well plates and then replated on 96-well plates (5,000 cells/well) in a volume of 100 µL and allowed to grow for 72 h. After experiments, 10 µL/well of WST-8 (cell counting kit 8, Sigma-Aldrich, 96992) reagent was added. After 3 h of incubation at 37 °C with 5% CO2, absorbance was read at 450 nm (Thermo, Multiscan Ascent). Medium without cells was used as a background control, subtracting this from the sample absorbance readings. Cell viability was calculated as a ratio of endpoint absorbance relative to control cells.

### Treatments

Heregulin β-1 (Sigma, H7660) was used at a working concentration of 20 ng.mL^-1^. For lysosome and proteasome inhibition, cells were treated for 4 h with 50 nM of bafilomycin A1 (Calbiochem, 196000) and 1 µM of bortezomib (Adooq Bioscience, A10160), respectively. Cycloheximide (Sigma, 01810) was used for translation inhibition at 25 µg.mL^-1^. For targeted inhibition of intracellular signaling proteins, trametinib (Adooq Bioscience, JTP-74057, [100] nM), rapamycin (Santa Cruz Biotechnology, sc-3504A, [100] nM), buparlisib (Adooq Bioscience, AT11016, [500] nM), selumetinib (Adooq Bioscience, A10257, [1] µM), SCH772984 (Adooq Bioscience, A12824, [1] µM) and MK-2206 (Adooq Bioscience, A10003, [2] µM) were used for the indicated time points.

### Immunofluorescence and confocal microscopy analyses

Cells were plated on µ-Slide 8-well dishes (Ibidi, 80826) and washed twice with ice-cold PBS before fixation in 4% paraformaldehyde for 10 min on ice. After fixation, cells were washed and then incubated with a permeabilization buffer (1X PBS/5% horse serum/0.3% Triton™ X-100) for 60 min. SiR-actin (Tebu-Bio, SC001) was used for counterstaining the cytoskeletal actin. Images were obtained using Zeiss LSM880 laser scanning confocal microscope. Airyscan images were processed with an Auto processing strength. Quantification of co-localization between SorLA and endosomal markers was performed using the ImageJ (Schneider et al., 2012) plugin ComDet (https://github.com/ekatrukha/ComDet) (Fréal et al., 2019).

### Dual-luciferase reporter assays

Firefly and Renilla luciferase activities of the same sample were sequentially measured with the results expressed as the ratio of firefly to Renilla luciferase activity. 3×10^4^ Cells were seeded per well of 48-well plates. Cells were co-transfected with 5’ SORL1 proximal promoter fragments in pGL4.10[luc2] vector (Promega, E6651) and pRL-TK Renilla control reporter vector (Promega, E2241) at relative DNA amounts of 95% and 5%, respectively. After the indicated treatments, cells were lysed and luciferase assays were performed using the Dual-Luciferase® Reporter Assay System (Promega, E1980) according to the manufacturer’s instructions. Luciferase activity was measured using Synergy H1 Hybrid Reader (BioTek). Each value of luciferase activity represents the mean of three internal replicates.

### Cell sorting

BT-474 cells were cultured on a monolayer of mCherry- or heregulin isoform 10 (SMDF)-overexpressing fibroblasts for 36 h. To bypass antibody-labeling steps prior to cell sorting, mCherry was overexpressed in BT-474 cells before starting the co-culture with SMDF-overexpressing fibroblasts. Before sorting, cells were detached using trypsin, harvested and put immediately on ice. Positive and negative selection of BT-474 cells was applied based on mCherry signal using Sony SH800S cell sorter. BT-474 cells were then pelleted and lysed in cell lysis buffer (CST, #9803) supplemented with 1% protease/phosphatase inhibitor cocktail (CST, #5872) prior to SDS-PAGE.

### Anchorage-independent 3D spheroid formation assays

An adapted protocol from (Peuhu et al., 2017) was used for spheroid growth assays. Cells were counted and equal amounts were plated in ultra-low attachment 96-well plate (Corning, CLS3474-24EA). Cells were allowed to proliferate for 7 days. Three internal replicates were plated for each sample. Images of spheroids were acquired using Eclipse Ti2 inverted microscope (Nikon) and spheroid volume was calculated using ImageJ (NIH).

### Matrigel-based multi-spheroid 3D growth assays

The bottom wells of a µ-Plate 96 well plate (ibidi, 89646) were filled with 10 µL of 50% matrigel (Corning, 356231) then the plate was centrifuged at 200 *g* for 20 min. The coated plate was incubated at 37 °C for 30 min. Wells were then filled with 20 µL of cell suspension (1000 cells, 1:1 ratio of BT-474 cells to TIFF) in 25% matrigel and incubated overnight at 37 °C. Wells were then filled with 65 µL of cell culture medium that was replaced every 2 days. Wells were imaged using the IncuCyte® S3 instrument (sartorius) and spheroid growth, reflected by mCherry fluorescence, was analyzed using the IncuCyte software (sartorious).

### Cloning for bimolecular fluorescence complementation (BiFC)

pDEST-SorLA-v1 and pDEST-SorLA- v2 were generated by first PCR amplifying the SorLA coding sequence from the plenti-SorLA-GFP vector (Pietilä et al., 2019) using the primers 5’- GGTACTCGAGGCCACCatggcgacacggagcagcaggaggga-3’ and 5’-GGTCGAATTCggctatcaccatggggacgtcatctgaaaatccag- 3’, and 5’-GGTACTCGAGGCCACCatggcgacacggagcagcagg- aggga-3’ and 5’-GGTCGAATTCggctatcaccatggggacgtcatctga- aaatccag-3’, respectively. PCR fragments were then subcloned into the pDEST-ORF-v1 (a kind gift from Darren Saunders, Addgene, #73637) (Croucher et al., 2016) or pDEST-ORF-v2 (Addgene, #73638) (Croucher et al., 2016) vectors using the XhoI/EcoRI restriction enzymes. For the pDEST-ERBB2-v2, pDEST-ERBB3-v1 and pDEST-ERBB3-v2 vectors, LR reactions (LR clonase II, ThermoFisher Scientific) were performed using the pDEST-ORF-v1 and pDEST-ORF-v2 destination vectors, and pENTR223-ERBB2 (a kind gift from William Hahn David Root, Addgene, #23888) (Johannessen et al., 2010) and pENTR223-ERBB3 (Addgene, #23874) (Johannessen et al., 2010) shuttle vectors. pENTR2b-mVenus was LR subcloned into pEF.DEST51 (ThermoFisher Scientific, #12285011) to generate the expression plasmid, pEF.DEST51-mVenus. All vectors were verified by analytical digests and sequencing.

### Lentivirus-mediated overexpression of Hrg β-1 and SMDF

Hrg β-1- and SMDF-coding sequences were LR sub-cloned from pENTR(tm)221 shuttle vectors (ThermoFisher Scientific Ultimate™ ORF, IOH80996, IOH80996) into pLenti7.3/V5-DEST (ThermoFisher Scientific, V53406) to generate expression plasmids. Lentiviral particles were generated in the 293FT packaging cell line (complete medium: high glucose DMEM, 10% FBS, 0.1 mM NEAA, 1 mM MEM Sodium Pyruvate, 6 mM L-Glutamine, 1% Pen/Strep and 0.5 mg/ml Geneticin) by transient transfection of transfer vector, either pLenti6.3/v5-DEST-SMDF (SMDF) or pLenti6.3/V5-DEST-Hrg β-1 (Hrg β-1), 2nd generation packaging plasmid-psPAX2 and envelope vector-pMD2 (kind gifts from Didier Trono, Addgene plasmids #12259 and #12260, respectively) with the ratio (7:2:1) using calcium-phosphate precipitation method (Graham and van der Eb, 1973). 72 h post transfection medium containing viral vectors was collected, concentrated for 2 h by ultracentrifugation in a swing-out rotor SW-32Ti (Beckman Coulter, Brea, US-CA) at 26000 *g*, resuspended in residual medium and flash frozen in liquid nitrogen. Functional titer of 1×10^8^ was measured in 293FT cells by FACS (BD LSRFortessa, Becton Dickinson). For stable overexpression, 8×10^4^ BT-474 cells and TIFF were seeded in a 24-well plate, 24 h later cells were transduced with MOI 1, 2 and 4 of Hrg β-1 or SMDF lentiviral stocks in low volume of full media. Medium containing viral particles was removed 16 h later. After 4 days, antibiotic selection was initiated with 35 µg/ml Blasticillin. Five days later, the selection was terminated. The MOI 4 condition with the highest (ca. 40%) survival rate was chosen for further experiments after validation of *NRG1* overexpression by qPCR.

### Zebrafish experiments

Zebrafish embryo experiments were carried out under license ESAVI/9339/04.10.07/2016 (National Animal Experimentation Board, Regional State Administrative Agency for Southern Finland). To test toxicity of tested drugs, 2 dpf zebrafish embryos of casper strain (White et al., 2008) were cultured in 96-well plates (1 embryo/well) and exposed to concentration series of tested drugs. All wells had a final concentration of 1% of DMSO and E3+PTU medium (5 mM NaCl, 0.17 mM KCl, 0.33 mM CaCl_2_, 0.33 mM MgSO_4_, 0.2 mM 4-phenylthiourea) at 33 °C. After 2 days of incubation, the mortality of embryos was evaluated under a stereomicroscope. The surviving embryos of each drug concentration were pooled, lysed for protein extraction and subsequently subjected to western blot analysis of biomarker proteins. For each drug, a lowest effective concentration resulting in robust decrease in biomarker was selected to be used in the following xenograft experiments.

Zebrafish embryo xenograft studies were essentially carried out as described in detail earlier (Paatero et al., 2018). In short, one day prior to transplantation GFP-MDA-MB-361 cells were transfected with control or SORL1 siRNAs. On the next day, the 2 dpf zebrafish embryos were immobilized in agarose, tumor cells suspended in PBS and injected into the brain from the dorsal side. One day post injection (1 dpi), successfully transplanted embryos were placed in CellView glass bottom 96-well plate (1 embryo/well) and drug treatments were initiated and embryos incubated in E3+PTU at 33°C. The xenografted embryos were imaged using a Nikon Eclipse Ti2 fluorescence microscope and a 2x Nikon Plan-Apochromat (NA 0.06) objective. Each embryo was imaged both at 1dpi and 4 dpi using brightfield illumination and a GFP fluorescence filter set (excitation with 470 nm LED). Each image was inspected manually to filter out severely malformed, dead or out of focus embryos. Next, the tumor area was measured using ImageJ (NIH). The fold change in tumor size was calculated as in Equation 1 (Eq. 1).

Eq 1. Fold change = (GFP intensity (4dpi))/(GFP intensity (1dpi))

### SPR analysis

SPR analysis was carried out by using a BIAcore3000 system (BIAcore, Uppsala). The SorLA ectodomain (Andersen et al., 2005) was immobilized on a CM5 chip at a density of 56 fmol/mm2. Subsequently, a concentration series of HER3 (ACRO Biosystems, ER3-H5223) or HER2 (ACRO Biosystems, HE2-H5212) were applied to the chip surface in 10 mM Hepes, pH 7.4/150 mM NaCl/5 mM CaCl2/0.005% Tween 20, and the respective BIAcore signals were expressed in RU corresponding to the difference in response between SorLA-coated and un-coated control flow channel. Kinetic parameters were determined by BIAEVALUATION 4.1 software.

### Statistical analyses

At least three biological replicates were performed for each experiment. The sample size (N) and the related statistical methods are described within figure legends. Significance was concluded when a probability value (P-value) was lower than 0.05. *NS: not significant; * p*<*0*.*05; ** p*<*0*.*01; *** 0*.*001; **** p*<*0*.*0001*.

## Acknowledgements

We thank J. Siivonen and P. Laasola for technical assistance. H. Hamidi is acknowledged for illustrations and editing the manuscript. The Ivaska laboratory is acknowledged for lively discussions and critical feedback on the manuscript. Guillaume Jacquemet, Jukka Westermarck, Majid Momeny, and Patrick Brest are acknowledged for critical reading of the manuscript. The Cell Imaging and Cytometry and Genome Editing core facilities (Turku Bioscience Centre, University of Turku and Åbo Akademi University and Biocenter Finland), the EuroBioimaging node in Turku and the Zebrafish core facility (Turku Bioscience Centre, University of Turku and Åbo Akademi University) are acknowledged for services, instrumentation, and expertise.

This study was supported by the Academy of Finland (J. Ivaska grant 312517), the Academy of Finland CoE for Translational Cancer Biology (J. Ivaska), the European Research Council CoG grant 615258 (J. Ivaska), the Sigrid Juselius Foundation (J. Ivaska) and the Finnish Cancer Organization (J. Ivaska). H. Al-Akhrass has been supported by the Finnish Cultural Foundation Central Fund 190150. J.R.W. Conway has been supported by the European Union’s Horizon 2020 research and innovation program under Marie Sklodowska-Curie grant agreement 841973. O.M. Andersen has been supported by The Independent Research Fund Denmark grant DFF-4004-00371. J. Kaivola has been supported by TuDMM Doctoral Program, University of Turku.

## Data availability

The authors declare that the data supporting the findings of this study are available within the article and from the authors on request.

## AUTHOR CONTRIBUTIONS

**Conception and design**: H. Al-Akhrass, J. Ivaska. **Development of methodology**: H. Al-Akhrass, J.R.W. Conway, A.S.A. Poulsen, J. Kaivola, A. Padzik, O.M. Andersen. **Acquisition of data**: H. Al-Akhrass, I. Paatero, O.M. Andersen. **Analysis and interpretation of data**: H. Al-Akhrass, J.R.W. Conway, I. Paatero, J. Kaivola, O.M. Andersen. **Writing**: H. Al-Akhrass, J. Ivaska. **Funding acquisition**: H. Al-Akhrass, J. Ivaska.

