## Supplementary Figures S1-5; Tables 1-3 for "A feed-forward loop between SorLA and HER3 determines heregulin response and neratinib resistance"

**Figure S1**

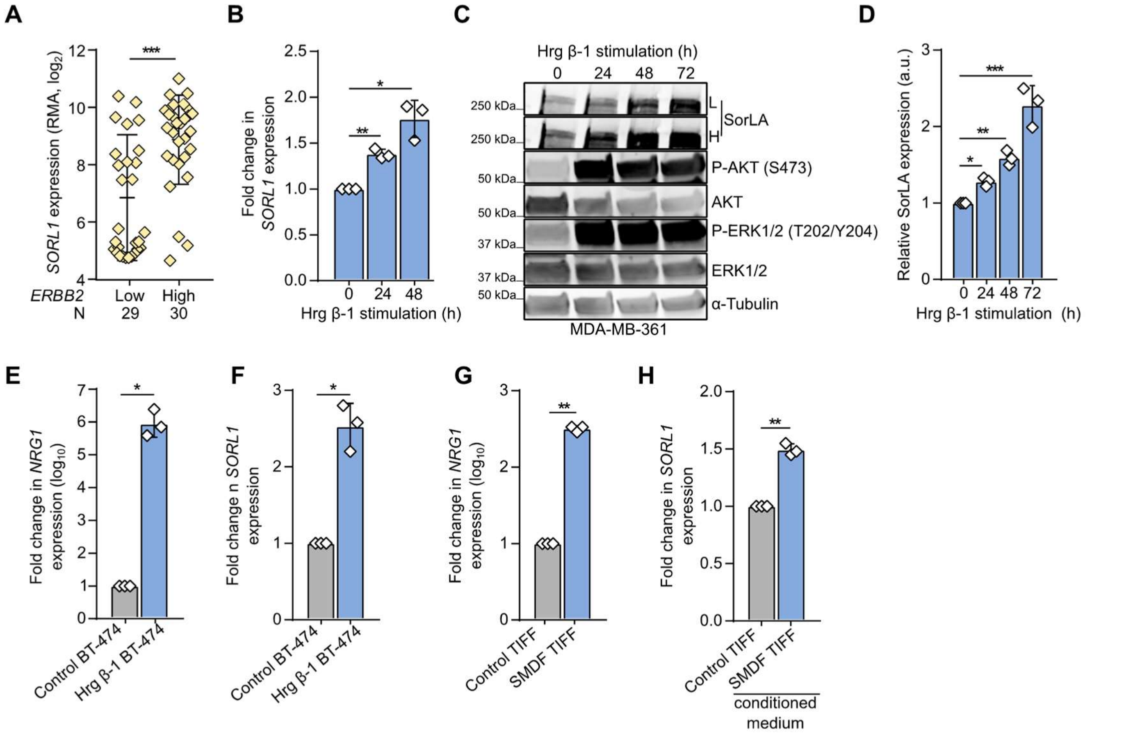


**Figure S1 related to Figure 1: HER3 signaling regulates *SORL1* expression**. **A**. *SORL1* expression is significantly higher in breast cancer cell lines with high *ERBB2* expression (CCLE; N=59). **B**. MDA-MB-361 cells were stimulated with 20 ng.mL$^{-1}$ Hrg β-1 for the indicated times. Quantification of *SORL1* mRNA levels, normalized to *HPRT1*, determined with RT-qPCR relative to non-stimulated (0 h) cells. **C**. MDA-MB-361 cells were stimulated with 20 ng.mL$^{-1}$ Hrg β-1 for the indicated times. Representative immunoblotting of SorLA, AKT(p)S473, total AKT, ERK1/2(p)T202/Y204, total ERK1/2, with α-tubulin as a loading control. **D**. Quantification of SorLA levels normalized to loading control and relative to non-stimulated (0 h) cells. **E&F**. Quantification of *NRG1* (**E**) or *SORL1* (**F**) mRNA levels, normalized to *HPRT1*, determined with RT-qPCR in mCherry control or Hrg β-1-expressing BT-474 cells relative to control cells. **G.** Quantification of *NRG1* mRNA levels, normalized to *HPRT1*, determined with RT-qPCR in mCherry control or SMDF-overexpressing fibroblasts (TIFF) relative to control cells. **H**. Quantification of *SORL1* mRNA levels, normalized to *HPRT1*, determined with RT-qPCR in BT-474 cells after 24 h of culture with conditioned media from either control or SMDF-TIFF relative to control TIFF.

Data are mean ± SD from three independent biological experiments; statistical analysis: Student's t-test (unpaired, two-tailed, unequal variance).

**Figure S2**

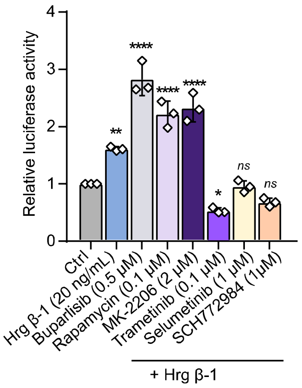


**Figure S2 related to Figure 2: heregulin-induced *SORL1* regulation requires HER3 signaling through ERK1/2**. Inhibitors of the ERK pathway decrease Hrg β-1-induced luciferase activity. P3-Luc was expressed in BT-474 cells together with pRL-TK Renilla luciferase transfection control and cells were treated with 20 ng.mL$^{-1}$ Hrg β-1 and the indicated inhibitors for 24 h. Luciferase activities are represented as fold increase from basal P3 luciferase activity. Data are mean ± SD from three independent biological experiments; statistical analysis: One-way ANOVA, Dunn's multiple comparisons test.

**Figure S3**

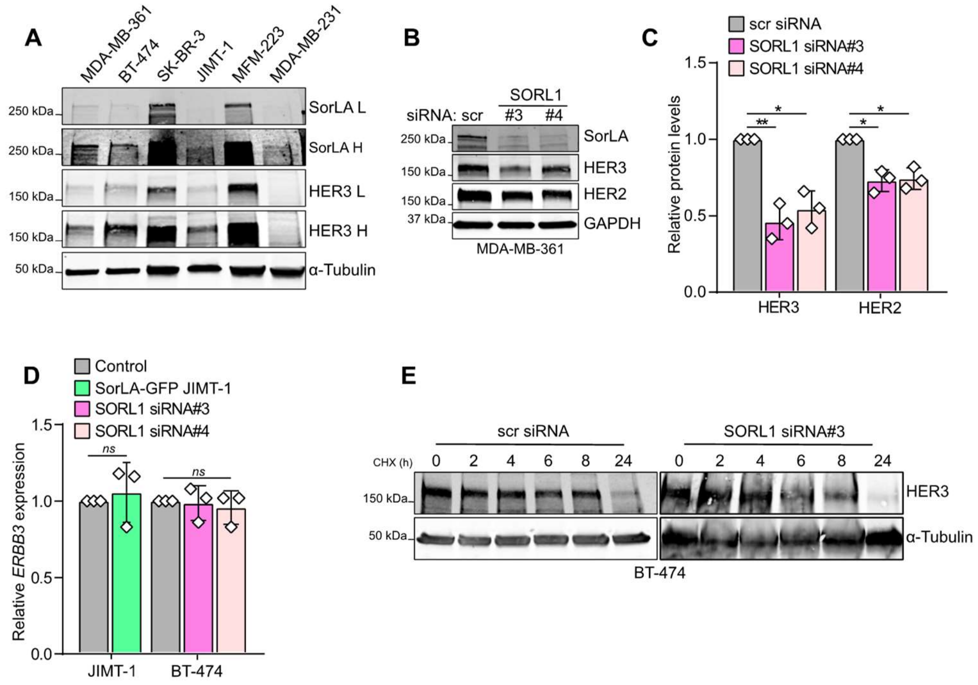


**Figure S3 related to Figure 3: SorLA regulates HER3 stability**. **A**. Representative immunoblotting of total HER3 and SorLA from the indicated breast cancer cell lines, with α-tubulin as a loading control. L and H indicate low and high contrast, respectively. **B**. Representative immunoblotting of HER2, HER3 and SorLA in control (Scr) and SorLA RNAi transfected MDA-MB-361 cells, with GAPDH as a loading control. **C**. Quantification of HER2 and HER3 levels normalized to loading control and relative to control-silenced cells. **D**. Quantification of *ERBB3* mRNA levels, normalized to *HPRT1*, determined with RT-qPCR in the cell lines transfected as indicated and relative to GFP-transfected controls (JIMT-1) or control siRNA-transfected BT-474 cells. **E**. Representative immunoblotting of HER3 in SorLA-silenced and control-silenced BT-474 cells treated with CHX for the indicated time points, with α-tubulin as a loading control. Double amount of protein was loaded for SorLA-silenced cells to allow equal detection of HER3.

Data are mean ± SD from three independent biological replicates. Statistical analyses: Student's t-test (unpaired, two-tailed, unequal variance). Scr = control non-targeting siRNA.

**Figure S4**

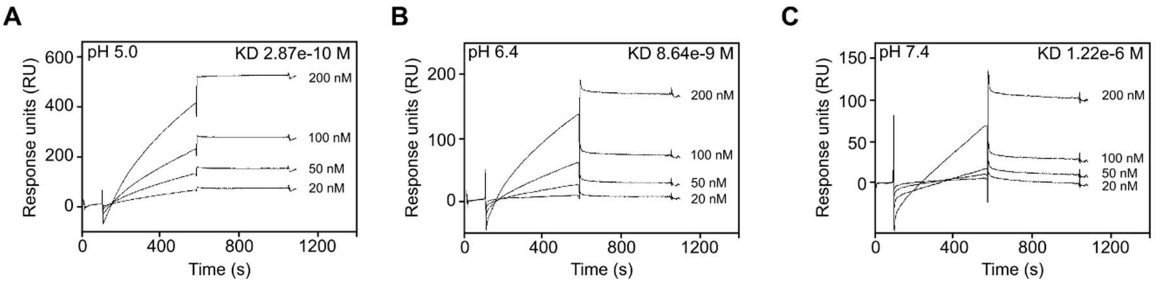


**Figure S4 related to Figure 4: SorLA interacts with HER2-HER3 dimers**. SorLA interacts with HER2 in a pH-dependent manner. SPR analysis of immobilized SorLA over a 20-200 nM concentration series of HER2 at pH 5.0 (**A**), 6.0 (**B**) and 7.4 (**C**).

**Figure S5**

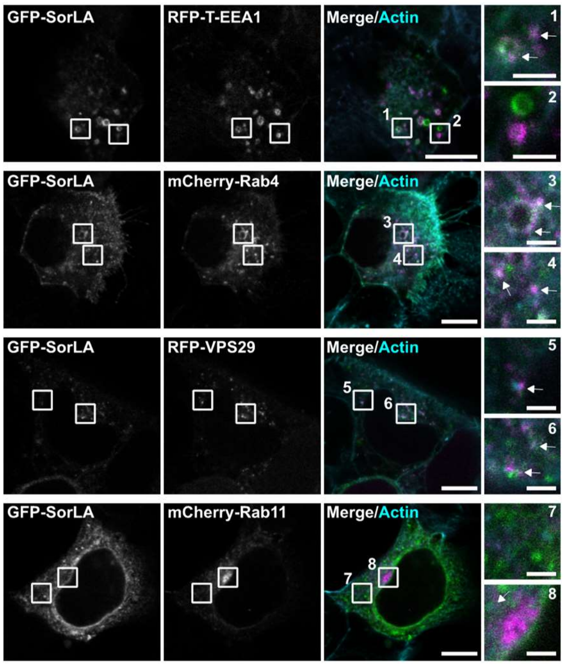


**Figure S5 related to Figure 5: SorLA regulation of HER2 and HER3 requires functional Rab4**. Representative confocal microscopy images of BT-474 cells co-expressing GFP-SorLA with the indicated endosomal markers. SiR-Actin was used for counterstaining the actin cytoskeleton. White arrows depict co-localizing signals. Scale bars: 10 µm. Scale bars (insets): 2 µm.

**Supplementary Table 1: kinetics of SorLA interaction with HER2 and HER3 calculated using BIAEVALUATION 4.1 software.**

| Receptors | pH | ka (1/Ms) | kd (1/s) | KD (M) |
|---|---|---|---|---|
| SorLA:HER2 | 5.0 | 2.135e4 | 6.126e-6 | 2.869e-10 |
| | 6.4 | 4.703e4 | 4.061e-4 | 8.635e-9 |
| | 7.4 | 1.866e3 | 2.284e-3 | 1.224e-6 |
| SorLA:HER3 | 5.0 | 7.088e4 | 2.871e-4 | 4.050e-9 |
| | 6.0 | 1.613e4 | 1.088e-4 | 6.745e-9 |
| | 7.4 | 2.160e5 | 2.075e-3 | 9.606e-9 |

**Supplementary Table 2: information related to the antibodies used in this study.**

| **Antibody** | **Manufacturer** | **Catalogue No.** | **Dilution** |
|---|---|---|---|
| HER3/ErbB3 (D22C5) | Cell Signaling Technology | #12708 | 1:1000 |
| p-AKT (S473) | Cell Signaling Technology | #9271S | 1:1000 |
| Total AKT | Cell Signaling Technology | #9272 | 1:1000 |
| phospho-ERK p-p44/42 MAPK T202/Y204 | Cell Signaling Technology | #4370S | 1 :1000 |
| Total ERK p44/p42 MAPK | Cell Signaling Technology | 9102S | 1:1000 |
| LR11 (SORL1) | BD Transduction Lab | 612633 | 1:1000 |
| HER2/ErbB2 (e2-4001 + 3B5) | Thermo Scientific | MA5-14057 | 1:1000 |
| GAPDH | HyTest | 5G4MaB6C5 | 1:2000 |
| α-tubulin | Hybridoma Bank | 12g10 | 1:5000 |
| β-actin | Sigma | A1978 | 1:5000 |

**Supplementary Table 3: list of previously published plasmids used in this study.**

| **Plasmid** | **Reference** |
|---|---|
| GFP-SorLA | (Pietilä et al., 2019) |

| | |
|---|---|
| RFP-T–EEA1 | (Navaroli et al., 2012) |
| RFP-VPS29 | (Hesketh et al., 2014) |
| mCherry-Rab4 | A gift from Michael Davidson (Addgene plasmid # 55125) |
| mCherry-Rab11 | A gift from Michael Davidson (Addgene plasmid # 55124) |
| GFP-Rab4$^{S22N}$ | (Arjonen et al., 2012) |
